# RAP2.3 is required for *MYB51* and *SIGMA3* expression during the response of *Arabidopsis thaliana* to multifactorial stress combination

**DOI:** 10.64898/2026.05.18.725943

**Authors:** Ranjita Sinha, María Ángeles Peláez-Vico, Devasantosh Mohanty, Lidia S. Pascual, Sara I. Zandalinas, Zhen Lyu, Ahmad Bereimipour, Rajeev K Azad, Trupti Joshi, Ron Mittler

## Abstract

In nature, plants are subjected to multiple environmental stress factors simultaneously or sequentially. Recent studies revealed that when three or more stress factors impact a plant simultaneously (termed ‘multifactorial stress combination’; MFSC), plant survival declines, even if the intensity of each individual stress involved in the MFSC is low. We previously identified RAP2.3 as a key transcription factor (TF) required for *Arabidopsis thaliana* survival, specifically under a MFSC of salt+excess light+heat stress (*i.e.,* S+EL+HS). Here we report that RAP2.3 is required for the expression of SIGMA3, a nuclear-encoded factor that directs plastid RNA polymerase to specific plastid promoters, and MYB51, a key stress response TF involved in glucosinolate metabolism and oxidative stress responses, specifically during a MFSC of S+EL+HS. Like *rap2.3* mutants, *myb51* and *sig3* mutants display significantly low survival rate specifically under the MFSC of S+EL+HS. Based on MYB51 gene regulatory network analysis and characterization of jasmonic acid (JA) mutants, we further reveal that suppression of JA signaling could play an important role in promoting plant survival under conditions of S+EL+HS. Our findings uncover an additional layer of the response of plants to MFSC, as well as identify potential targets for breeding crops with enhanced tolerance to climate change.

## 1. INTRODUCTION

In nature, plants and other organisms are continuously subjected to a multitude of different environmental factors that change in their intensity, duration, and/or frequency of occurrence. Some of these environmental conditions could become stressors if they pass a certain threshold resulting in the activation of different acclimation and defense mechanisms in plants. As multiple stressors may impact a plant simultaneously or sequentially, plants growing in nature are often subjected to conditions of ‘stress combination’ (*i.e.,* two or more stressors impacting a plant simultaneously or sequentially; Mittler 2006; Speißer et el. 2022; Zandalinas and Mittler 2022; Zhou et al. 2023; Alptekin and Kunkowska 2024; Sato et al. 2024; Rongstock et al. 2026). Examples for these include nutrient deprivation combined with water deficit, heat stress combined with water deficit and/or high salinity, and more (*e.g.,* Rizhsky et al. 2004; Balfagón et al. 2024; Pardo-Hernández et al. 2024; Senizza et al. 2024; Raza et al. 2025; Johnson et al. 2026). Multiple studies revealed that the molecular, metabolic and physiological responses of plants to conditions of stress combination are unique and cannot be inferred from studying the response of plants to each of the different stresses, involved in the stress combination, applied individually (summarized in Mittler 2006; Zandalinas and Mittler 2022; Zhou et al. 2023; Balfagón et al. 2024; Sato et al. 2024; Senizza et al. 2024; Raza et al. 2025; Rongstock et al. 2026; Johnson et al. 2026). Thus, the ‘reductionist approach’ of subjecting a plant to a single stress condition (while controlling all other conditions) may not accurately reveal how plants respond to stress under conditions of stress combination, that mostly occur in nature.

Recent studies of plant responses to a combination of two different stresses occurring simultaneously identified different transcription factors (TFs) such as WRKY48 (heat and excess light combination; Balfagón et al. 2024) and NIN-LIKE PROTEIN 7 (NLP7; nutrient deficiency and water deficit combination; Johnson et al. 2026), different regulatory processes such as OPEN STOMATA 1 (OST1) inactivating TARGET OF TEMPERATURE 3 (TOT3; heat and water deficit combination; Xu et al. 2025), as well as the physiological process of ‘differential transpiration’ (between reproductive and vegetative tissues during a combination of water deficit and heat; Sinha et al. 2022; 2023; 2025a; 2025b), as uniquely activated during stress combination. In addition to these studies of 2-stress combination, recent studies of multiple (3 or more) stress combinations in plants, termed ‘multifactorial stress combination’ (MFSC) uncovered a new principle in biological sciences, termed the ‘MFSC principle’. Multifactorial stress combination is a type of stress combination defined as three or more stressors impacting a plant simultaneously or sequentially (Zandalinas et al. 2021a; 2024; Zandalinas and Mittler 2022). In studies in which plants, such as Arabidopsis (*Arabidopsis thaliana*; Zandalinas et al. 2021b), tomato (*Solanum lycopersicum*; Pascual et al. 2023; 2025a), soybean (*Glycine max*; Peláez-Vico et al. 2024), rice (*Oryza sativa*; Sinha et al. 2024), or corn (*Zea mays*; Sinha et al. 2024) were subjected to 4-6 different abiotic stressors simultaneously (in most or all possible combinations) it was found that with the increase in stress complexity (*i.e.,* the number of stresses simultaneously impacting a plant), plant physiological performance, growth, and even survival, dramatically declined, even if the level of each stress involved in the MFSC did not have a significant impact, or had a minimal effect, on plants (defined as the ‘MFSC principle’). A similar effect was also found when *Chlamydomonas reinhardtii* cells were subjected to MFSC (Pascual et al. 2025b). As environmental conditions on our planet are becoming more extreme due to climate change and increased pollution levels (Rillig et al. 2019; Lee et al., 2023; Richardson et al. 2023; Zhou et al. 2023), more studies of MFSC are needed, so that we could identify MFSC-specific genes and pathways that will help us develop stress combination/MFSC resilient crops (Rivero et al. 2022; Zandalinas et al. 2024).

We recently reported that the TF bHLH35 (At5g57150) is specifically required for the survival of Arabidopsis plants during a MFSC of salinity (S), excess light (EL), and heat (HS) stresses (*i.e.,* S+EL+HS), but not when each of these stresses are applied individually or in any other 2-stress combination (Sinha et al. 2025c). During the S+EL+HS MFSC, bHLH35 was found to interact with NAC069 and regulates the expression of LBD31 (At4g00210). In addition, bHLH35 was found to be required for RAP2.3 (At3g16770) expression during the MFSC of S+EL+HS (Sinha et al. 2025c). While we found that, through activating LBD31, bHLH35 regulates flavonoids metabolism that suppresses reactive oxygen species (ROS) accumulation, the role of RAP2.3 during MFSC remained unclear (Sinha et al. 2025c). Here, we report on the characterization of two independent mutant alleles for RAP2.3 (*rap2.3_1* and *rap2.3_2*). Both alleles express constitutively high levels of the *RAP2.3* transcripts under controlled growth conditions, as well as accumulate high levels of *RAP2.3* transcripts, compared to wild type (WT), in response to the MFSC of S+EL+HS. In addition, like the *bhlh35* mutants (Sinha et al. 2025c), when subjected to a MFSC of 5 different abiotic stresses in increasing order of complexity, both *rap2.3* mutants were found to be deficient in survival, specifically under conditions of S+EL+HS. Using transcriptomics and gene regulatory network (GRN) analyses of WT and one of the RAP2.3 mutants (*rap2.3_1*), we further identified MYB51 (At1g18570) and SIGMA3 (At3g53920) as two transcriptional regulators that require RAP2.3 function for their expression under conditions of S+EL+HS. Interestingly, two independent mutant alleles deficient in the expression of each of these two transcriptional regulators were also found to be deficient in plant survival specifically under conditions of S+EL+HS. Based on MYB51 GRN analysis and jasmonic acid (JA) mutants characterization, we further reveal that suppression of JA signaling could play an important role in promoting plant survival under conditions of MFSC. Taken together, our findings reveal that constitutive expression of RAP2.3 disrupts the response of plants to MFSC, and that this disruption could be linked to the function of MYB51 and SIGMA3, as well as to JA signaling.

## 2. MATERIALS AND METHODS

### 2.1. Plant growth, stress treatments, and mutant screening

*Arabidopsis thaliana* wild-type Col-0 (WT) and two independent T-DNA insertion mutants of RAP2.3/ERF7 were tested for survival under the following individual treatments and their different combinations on plates containing ½ Murashige and Skoog (½ MS; Caisson Labs, UT, USA; MSP09) medium, pH 5.8, with 1% phytoblend (Caisson Labs, UT, USA; PTP01): CT (control; ½ MS, 21 °C, 50 µmol m^-2^ s^-1^, pH 5.8), Cd (cadmium; ½ MS, 21 °C, 50 µmol m^-2^ s^-1^, pH 5.8, 5 µM CdCl_2_), EL (excess light; ½ MS, 21 °C, pH 5.8, 700 µmol m^-2^ s^-1^), HS (heat stress; ½ MS, 50 µmol m^-2^ s^-1^, pH 5.8, 33 °C), S (salt stress; ½ MS, 21 °C, 50 µmol m^-2^ s^-1^, pH 5.8, 50 mM NaCl), and PQ (paraquat; ½ MS, 21 °C, 50 µmol m^-2^ s^-1^, pH 5.8, 0.05 µM paraquat) as described in (Zandalinas et al. 2021b; Sinha et al. 2025c; Table S1). All other mutants were tested for survival only under CT conditions, individual and all possible combinations of S, EL and HS as described above. All the T-DNA insertion mutants used in the study were obtained from Arabidopsis Biological Resource Center (ABRC; https://abrc.osu.edu/researchers; Table S2).

### 2.2. RNA-Seq analysis

*A. thaliana* wild-type Col-0 and *rap2.3_1* seedlings (about 125-150) were grown horizontally for 6 days on separate sets of ½ MS plates with and without salt (50 mM NaCl) in three biological replicates. Six-day old seedlings were then subjected to the individual and combined stresses of S, EL (1.5 hours) and HS (1.5 hours) as described above. Whole seedlings were sampled for each stress treatment and control in three replicates as described by Zandalinas et al. (2021b) and Sinha et al. (2025c). Total RNA was isolated using RNAeasy plant mini kit (Qiagen, MD, USA; 74904). RNA libraries for sequencing were prepared by Novogene Co. Ltd (https://en.novogene.com/website, Sacramento, CA) using standard Illumina protocols. RNA sequencing was performed using NovaSeq 6000 PE150 by Novogene Co. Ltd.

Quality control, read trimming, read alignment, transcript assembly, quantification, and differential expression analysis of RNAseq data were performed as explained previously (Sinha et al. 2025c). KEGG enrichment and overrepresented gene ontology (GO) terms (*P* < 0.05) were conducted using g:Profiler (https://biit.cs.ut.ee/gprofiler/gost; Kolberg et al. 2023). Heatmaps were generated using Morpheus (https://software.broadinstitute.org/morpheus/) and Venn diagrams were created in VENNY 2.1 (BioinfoGP, CNB-CSIC; https://bioinfogp.cnb.csic.es/tools/venny/index.html). UpSet plots were generated using the UpSetR (Conway et al. 2017) package. RNA-seq data files are available at GEO (GSE330879).

### 2.3. Promoter motif analysis

The promoter sequences 3000 bp upstream of the start site were downloaded from The Arabidopsis Information Resource (TAIR; https://www.arabidopsis.org/) for genes in Class I, Class II and genes coding for JA biosynthesis pathway enzymes. PBM and DAP motif files for RAP2.3 and MYB51, respectively, were downloaded from Plant Transcription Factor Database (PlantTFDB; https://planttfdb.gao-lab.org/). Presence of binding motif in the promoters was performed in MEME Suite 5.5.4 (https://meme-suite.org/meme/index.html) using the FIMO motif scanning tool (https://meme-suite.org/meme/tools/fimo; Grant et al. 2011).

### 2.4. Quantitative real-time PCR

Total RNA for quantitative real-time PCR (qRT-PCR) analysis was isolated from Arabidopsis seedlings of WT, *rap2.3* mutants and over-expression transgenics growing under CT and stress conditions using the stress and RNA isolation protocols described above. Details of the primer pair used for the real-time PCR analysis are in Table S3.

### 2.5. Gene regulatory network analyses

Gene Regulatory Network analysis was performed using DIANE (Differential Inference Analysis of Expression (Cassan et al. 2021), as described in Sinha et al., (2025c). Please see Supplementary text for more details.

### 2.6 *RAP2.3* over-expression line

*A. thaliana* Col-0 WT plants were transformed with the RAP2.3 CDS driven by the 35S-CaMV promoter. The coding sequence of RAP2.3 was cloned into pCAMBIA1304 binary vector downstream of the 35S-CaMV promoter and upstream of the mGFP5 reporter gene. Cloning was performed using In-Fusion cloning kit (Takara, CA, USA; 638955). After confirmation by sequencing, the recombinant vector was transferred to *Agrobacterium tumefaciens* GV3101. *A. thaliana* plants were transformed by floral dip. T1 seeds were screened on ½ MS media supplemented with 12 µg mL^-1^ hygromycin as described in Sinha et al. (2025c). Three independent homozygous transgenic lines were generated (T3) and tested for survival under conditions of stress combination as described above. Over-expression was confirmed by qRT-PCR. Details of the primer pair used for cloning are in Table S3. Nitric oxide imaging of WT and RAP2.3 overexpressing plants was conducted using 10-day-old seedlings grown on plates under controlled growth conditions according to Mohanty et al., (2025).

### 2.7 Data analysis

Two-way ANOVA with Fisher’s LSD post hoc tests were used to test significant differences between the percent survival of WT and the different mutant genotypes. The mean percent survival difference at a *P* < 0.05 was considered as significant. One-way ANOVA with Fisher’s LSD post hoc tests was used to test for significant differences between the gene expression (RQ) of WT and the different genotypes. Unpaired t-test with Welch’s correction was used to determine significant differences between the expression of *RAP2.3* in WT plants subjected to CT or S+EL+HS.

## 3. RESULTS

### 3.1. Characterization of *rap2.3* mutants subjected to a 5-stress MFSC

We previously reported that, compared to WT, two independent mutant alleles of the *Rap2.3* gene (*rap2.3_1*, SALK_054382 and *rap2.3_2*, CS470425) are more susceptible to a MFSC of S+EL+HS (Sinha et al. 2025c). However, whether RAP2.3 is required for survival under additional MFSC conditions, and what is the nature of the *rap2.3_1* and *rap2.3_2* mutations in the *Rap2.3* gene (see below) remained unknown. To address these questions we subjected WT and the two *rap2.3* mutants to a MFSC of five different low level abiotic stress conditions [*i.e.,* salt (S; 50 mM NaCl), EL (700 μmol m^−2^ s^−1^), HS (33°C), the herbicide paraquat (PQ; 0.05 μM), and/or the heavy-metal cadmium (Cd; 5 μM CdCl_2_), Table S1] in all possible combinations and compared their survival to that of WT. As shown in Figure 1A, even at this complex level of 5-stress MFSC, compared to WT, the two *rap2.3* mutants were specifically sensitive to the combination of S+EL+HS. Thus, compared to WT, the two *rap2.3* mutants had a significantly lower survival rate under conditions of S+EL+HS (Figure 1A). In addition to S+EL+HS, the two *rap2.3* mutants were also sensitive to other (higher order) MFSC that included S+EL+HS (*i.e.,* S+PQ+EL+HS, S+Cd+EL+HS, and S+PQ+Cd+EL+HS), but then so was WT. These results suggested that the addition of Cd and/or PQ to the S+EL+HS MFSC resulted in conditions that impaired the survival of both WT and the *rap2.3* mutants. Interestingly, the combination of PQ+Cd+EL+HS did not impact the survival or WT or the *rap2.3* mutants, highlighting the key role of S added to EL+HS in enhancing the susceptibility of *rap2.3* mutants under conditions of S+EL+HS. Since both Cd and PQ stresses are likely to result in enhanced ROS levels in cells (Chmielowska-Bąk et al. 2014; Mittler et al. 2022; Niekerk et al. 2024), it is possible that the MFSC of S+EL+HS renders plant especially susceptible to oxidative stress. This possibility is supported by our previous findings that a MFSC of S+EL+HS causes an enhanced accumulation of ROS in plants (Zandalinas et al. 2021b; Sinha et al. 2025c) and that, under conditions of S+EL+HS, *bhlh35* mutants accumulate high levels of ROS that, once suppressed by the addition of naringenin (a flavonoid), restored their survival (Sinha et al. 2025c). It should however be noted that the addition of naringenin did not restore the survival of the *rap2.3_1* and *rap2.3_2* mutants under conditions of S+EL+HS (Sinha et al. 2025c). Thus, in contrast to WT or the *bhlh35* mutants, the cause of *rap2.3* mutants’ lethality under conditions of S+EL+HS may not be associated with enhanced ROS levels.

**Figure 1.**
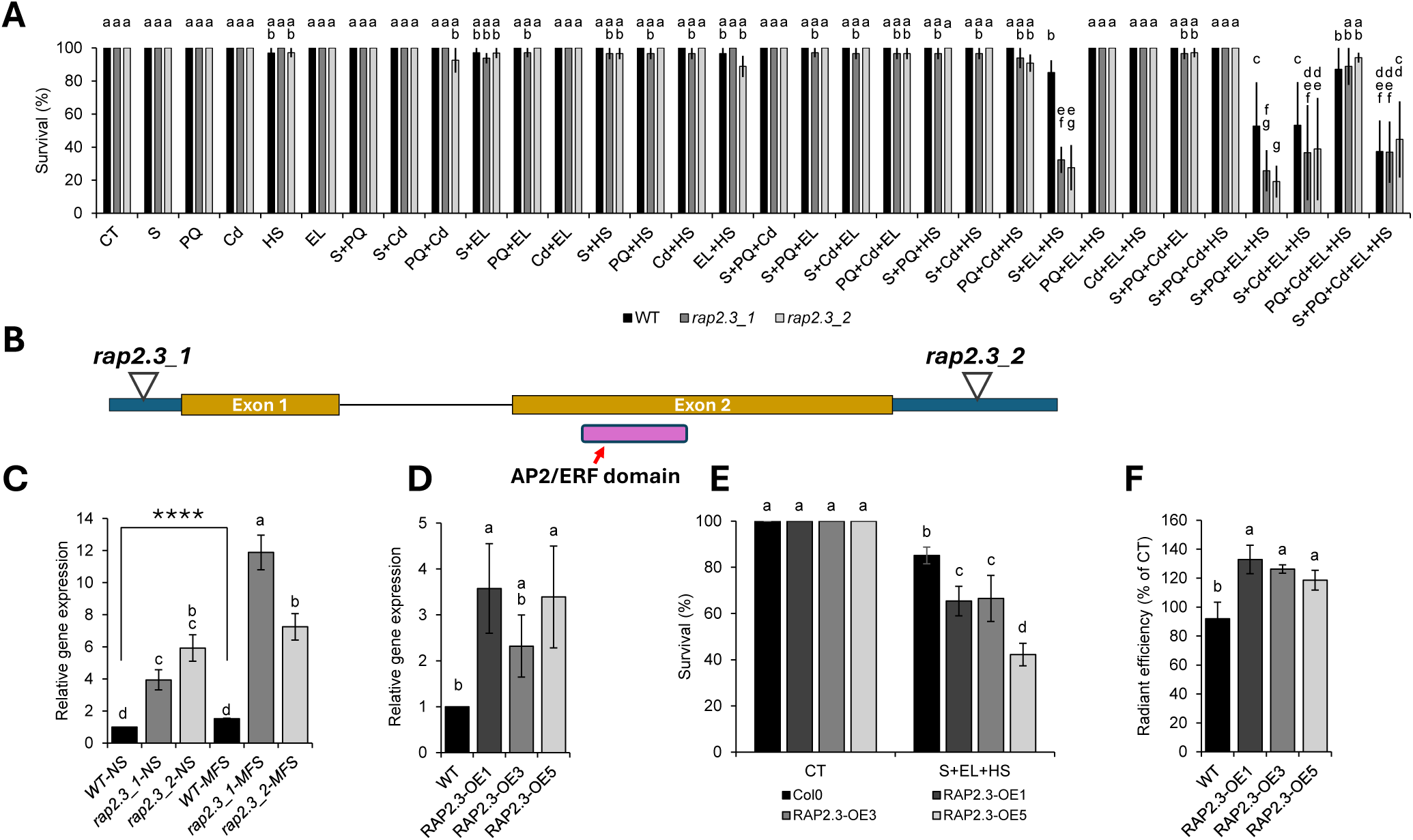
Characterization of *rap2.3* mutants. **A.** Survival of wild type (WT) and *rap2.3* (*rap2.3_1*, SALK_054382 and *rap2.3_2*, CS470425) seedlings grown under controlled growth conditions (CT), or subjected to a multifactorial stress combination (MFSC) of five different low level abiotic stress conditions [*i.e.,* salt (S; 50 mM NaCl), excess light (EL; 700 μmol m^−2^ s^−1^), heat stress (HS; 33°C), the herbicide paraquat (PQ; 0.05 μM), and/or the heavy-metal cadmium (Cd; 5 μM CdCl_2_)] in all possible combinations. **B.** Schematic presentation of the *Rap2.3* gene with position of the two T-DNA inserts for *rap2.3_1* and *rap2.3*_2 indicated. **C.** Quantitative real-time PCR (qRT-PCR) analysis of *RAP2.3* transcripts steady-state levels in WT and *rap2.3* mutants grown under CT (NS) conditions or subjected to a MFSC of S+EL+HS (MFS). **D.** qRT-PCR analysis of RAP2.3 transcripts steady-state levels in WT plants or transgenic plants that overexpress RAP2.3 (RAP2.3-OE) grown under CT conditions. **E.** Survival of WT plants and RAP2.3-OE grown under CT conditions or subjected to a MFSC of S+EL+HS. **F.** Nitric oxide (NO) accumulation in WT and RAP2.3-OE plants grown under CT conditions. Two-way ANOVA followed by Fisher’s LSD post hoc test was used to determine significant differences between means in panels A and C and one-way ANOVA followed by Fisher’s LSD post hoc test was used to determine significant differences between means in panels D-F. Different letters denote statistically significant differences; *P* ≤ 0.05. Unpaired t-test with Welch’s correction was used to determine significant differences between means of qRT-PCR of WT-NS and WT-S (****, *P* ≤ 0.0001) in panel C. Abbreviations: AP2, APETALA 2; Cd, cadmium; CT, control; EL, excess light; ERF, ethylene response factor; HS, heat stress; MFS, S+EL+HS; NS, control condition; OE, over-expression lines; PQ, paraquat; S, salt; WT, wild type Col0.

### 3.2. *Arabidopsis thaliana* plants with elevated levels of *RAP2.3* expression are susceptible to a MFSC of S+EL+HS

The T-DNA insertions in the two *rap2.3* mutants are found at the 5’ or 3’ untranslated regions of the *rap2.3* gene (Figure 1B). To determine how these insertions impacted the steady-state level of the *RAP2.3* transcript under control or S+EL+HS conditions, we measured its level in WT and the *rap2.3* mutants using quantitative real-time PCR (qRT-PCR) analysis. As shown in Figure 1C, application of S+EL+HS to WT plants resulted in a significantly elevated expression of *RAP2.3* (in agreement with Zandalinas et al. 2021b and Sinha et al. 2025c; Unpaired t-test with Welch’s correction was used to determine significant differences between control and S+EL+HS of WT plants; *p* ≤ 0.0001). Interestingly, the basal steady-state level of transcripts encoding RAP2.3 in the two *rap2.3* mutants was higher than that of control WT plants (as well as higher than that of S+EL+HS-stressed WT plants), and the expression level of *RAP2.3* transcripts was further elevated in response to S+EL+HS conditions (to levels that were statistically higher than that of WT in response to S+EL+HS; Figure 1C; Two-way ANOVA followed by a Fisher LSC post hoc test; *P* ≤ 0.05). These results suggest that the elevated basal expression levels of *RAP2.3* in the *rap2.3* mutants could be responsible for the observed phenotype shown in Figure 1A. To test this possibility, we generated transgenic plants that express the *RAP2.3* transcript under the control of the 35S-CaMV promoter (*RAP2.3-OE*) and tested their survival under conditions of S+EL+HS. As shown in Figure 1D, these plants displayed high levels of *RAP2.3* transcript expression when grown under controlled growth conditions. In addition, compared to WT, these plants were more susceptible to the MFSC of S+EL+HS (Figure 1E). As RAP2.3 is typically subjected to protein degradation by the N-degron pathway under aerobic conditions (Licausi et al. 2011; Gibbs et al. 2011; 2014), our findings that elevated expression levels of the *RAP2.3* transcript in cells result in a specific S+EL+HS MFSC sensitivity phenotype (Figure 1), could suggest that the high expression level of RAP2.3 can overcome the degradation of the RAP2.3 protein. This possibility is supported by suppressed expression of different members of the N-degron pathway in the *rap2.3* mutant under conditions of S+EL+HS (Figure S1). In addition, in accordance with RAP2.3 role in nitric oxide (NO) signaling (León et al., 2020), compared to WT plants, plants that overexpressed RAP2.3 were found to accumulate altered levels of NO (Figure 1F), providing further support to the possibility that overexpression of RAP2.3 can overcome the N-degron pathway. Taken together, the findings presented in Figure 1 could suggest that constitutive expression of *RAP2.3* in the *rap2.3* mutants or *RAP2.3-OE* transgenic plants results in sufficient accumulation of RAP2.3 protein that either suppresses pathways that are essential for acclimation to S+EL+HS, or activates pathways that conflict with S+EL+HS acclimation. The possibility that the N-degron pathway is overwhelmed in *rap2.3* mutants could be addressed in future studies by testing the survival phenotype of transgenic plants (WT) with constitutive overexpressing an N-degron-resistant variant of RAP2.3 (Gibbs et al. 2014; León et al. 2020). These are expected to yield a similar or even stronger phenotype to the current *rap2.3* or the *RAP2.3-OE* plants we have already characterized (Figure 1).

### 3.3. Transcriptomics analysis of WT and *rap2.3_1* during MFSC

To determine the role of RAP2.3 during MFSC, we conducted RNA-Seq analysis of WT and one of the RAP2.3 mutants (*rap2.3_1*) subjected to S, EL, and HS in all possible combinations (Tables S4-S5). As shown in Figure 2A, a read map alignment analysis of the *Rap2.3* gene under CT, and S, EL, and/or HS in all possible combinations supported the qRT-PCR analysis shown in Figure 1C and revealed that transcripts encoding *RAP2.3* are elevated in their expression under control and all stress conditions in the *rap2.3_1* mutant. This analysis also revealed that the only intron found in the *Rap2.3* gene is efficiently spliced under all growth conditions, with some minor exceptions during HS, HS+EL, and S+EL+HS (Figure 1C). To identify transcripts significantly altered in their expression (up or down) in WT or *rap2.3_*1, in response to all possible combinations of S, EL, and HS (*i.e.,* S, EL, HS, S+EL, S+HS, EL+HS, S+EL+HS), we conducted a differential transcript expression analysis between control (CT) and each different stress treatment, for WT and the *rap2.3_1* mutant (Figure 2B; Tables S4-S6). This analysis revealed that over 2800 transcripts are altered in WT or the *rap2.3_*1 mutant in response to each of the different stresses or their combinations (Figure 2B). Venn diagrams comparing the transcripts significantly altered in their expression in WT or the *rap2.3_1* mutant under each of the different stresses studied identified multiple transcripts that are altered in their expression in WT plants but not the *rap2.3_1* mutant (indicated by arrows; putative RAP2.3-dependent; Figure 2B). An UpSet plot of the overlap between these different groups of transcripts (specifically altered in WT but not the *rap2.3_1* mutant in response to each of the different stress treatments) revealed that there was no overlap between the different transcripts that depend on RAP2.3 for their expression during the different stresses and their combinations (Figure 2B). Similarly, there was no overlap between the transcripts significantly altered in the *rap2.3_1* mutant, but not in WT, in response to all stresses and their combinations studied (Figure S2). These results suggest that RAP2.3 could have a unique regulatory function under each different stress condition or stress combination. A similar finding was previously reported for bHLH35 that functions upstream to RAP2.3 under the different stress conditions tested (Sinha et al. 2025c), providing further support to the link between these two TFs (*i.e.,* bHLH35 and RAP2.3).

**Figure 2.**
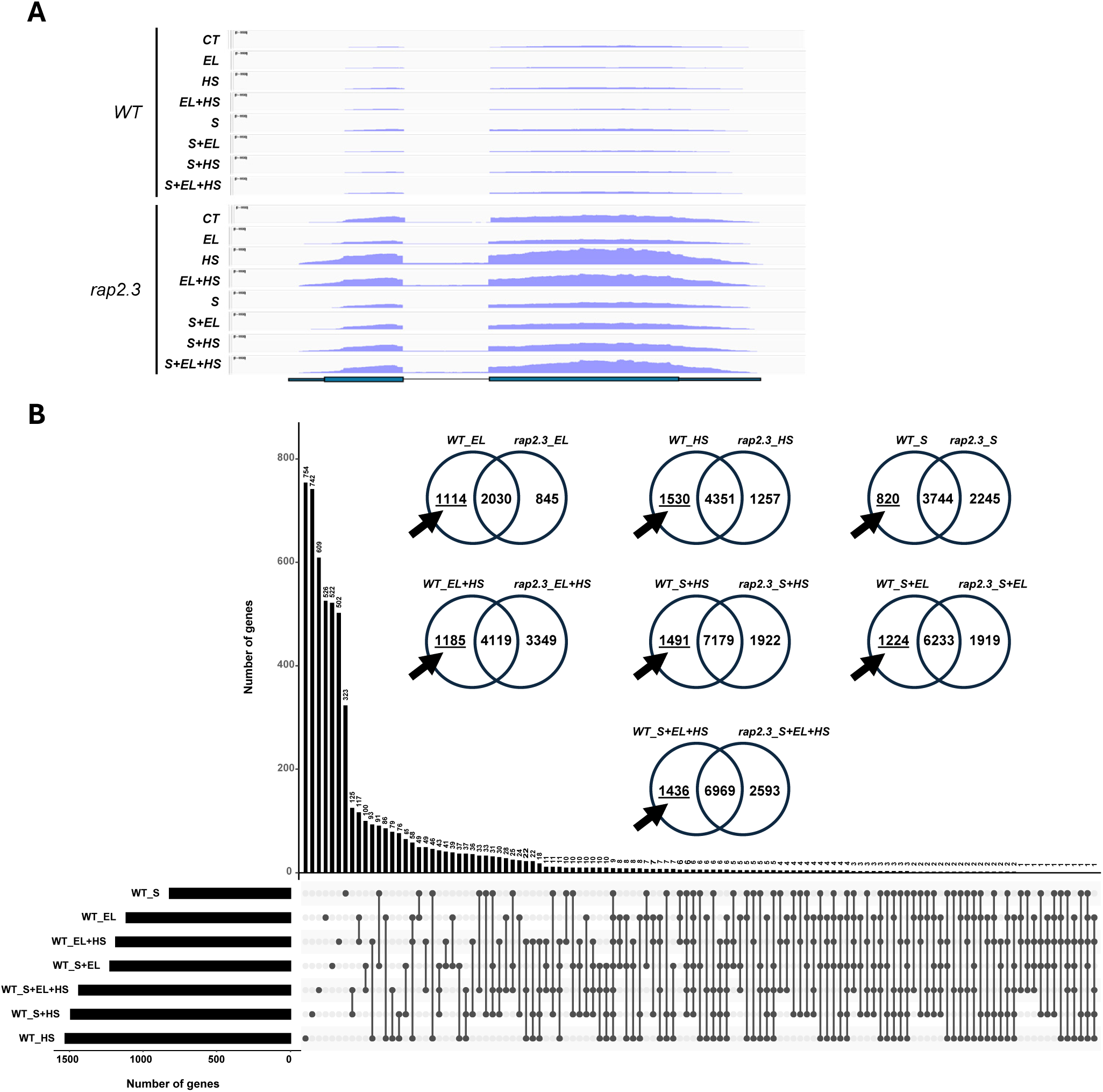
Transcriptomics analysis of wild type and *rap2.3_1* seedlings subjected to salt, excess light and heat stresses in all possible combinations. **A.** RNA-Seq read maps aligned to the *RAP2.3* gene in wild type (WT) and *rap2.3_1* seedlings grown under controlled growth conditions (CT), or subjected to subjected to salt (S), excess light (EL), and/or heat stress (HS) conditions in all possible combinations. **B.** An UpSet plot and Venn diagrams showing the overlap between transcripts significantly altered in their expression in WT or the *rap2.3_1* mutant in response to all stress treatments compared to CT. Arrows indicate transcripts with significantly altered expression in WT, but not the *rap2.3_1* mutant. Negative binomial Wald test followed by Benjamini-Hochberg correction (*P* ≤ 0.05) was used for statistical significance between transcript expressions. Abbreviations: CT, control; EL, excess light; HS, heat stress; S, salt; WT, wild type Col0.

### 3.4. Comparative transcriptomics analysis of WT and *rap2.3_1* under conditions of S+EL+HS

As the function of RAP2.3 appears to be specific for each stress/stress combination condition studied (Figures 2B, S2), we focused our analysis on the MFSC of S+EL+HS, that resulted in the specific mortality rate phenotype of the *rap2.3* mutants (Figure 1A). Thus, we conducted a GRN analysis of transcripts significantly altered in WT plants in response to S+EL+HS (8405 transcripts; Figure 2B) with a focus on RAP2.3. As shown in Figure 3A, our GRN analysis identified 312 transcripts associated with RAP2.3 function during a MFSC of S+EL+HS. These were classified by the GRN analysis into 8 different clusters (Table S7). A GO analysis of these clusters revealed that they are enriched in transcripts involved in flavone metabolism, glucosinolate biosynthesis, abiotic stress responses, epigenetic regulation, and ethylene signaling (Figure 3B; Table S8). A link between RAP2.3 and ethylene signaling was previously reported (Sinha et al. 2025c), supporting our GRN analysis.

**Figure 3.**
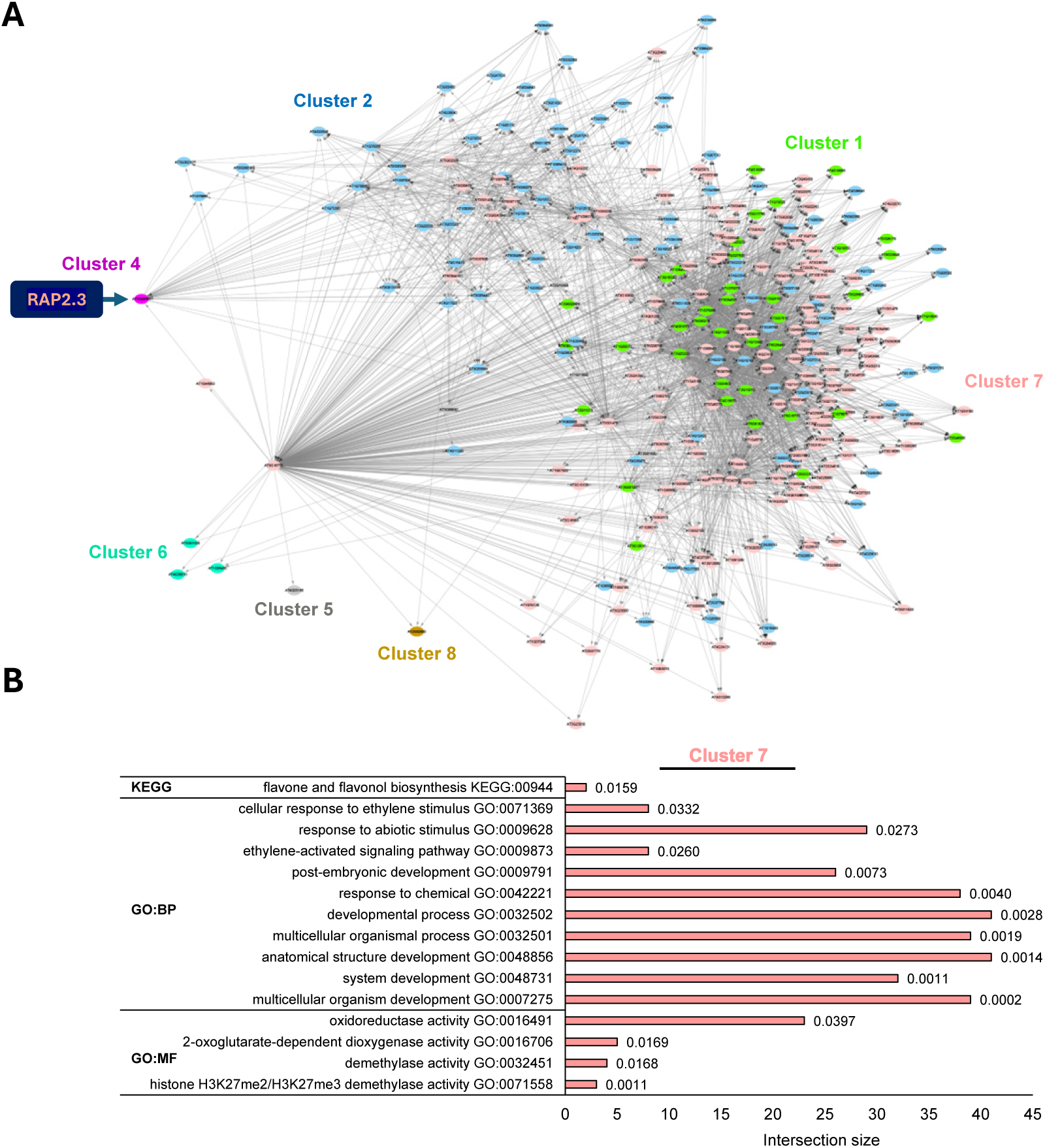
Gene regulatory network analysis of *RAP2.3* during multifactorial stress combination. **A.** Gene regulatory network (GRN; Differential inference analysis of expression; DIANE) analysis of *RAP2.3* during a multifactorial stress combination (MFSC) of salt (S), excess light (EL), and/or heat stress (HS). **B.** Gene ontology (GO) analysis of transcripts associated with *RAP2.3* function during a MFSC of S+EL+HS. Abbreviations: BP, biological process; GO, gene ontology; MF, molecular function; KEGG, Kyoto Encyclopedia of Genes and Genomes.

We further divided the transcripts significantly altered in their expression in WT and *rap2.3_1* into Class I (transcripts altered in their expression in WT but not in *rap2.3_1* in response to S+EL+HS; 1436 transcripts) and Class II (transcripts altered in their expression in *rap2.3_1* but not in WT in response to S+EL+HS; 2593 transcripts; Figure 4A). GO and KEGG analyses of these two groups identified multiple abiotic and biotic stress-response transcripts in Class I, and multiple abiotic stress response, as well as proteosome-, 2-oxocarboxylic acid metabolism-, and amino acid biosynthesis-related transcripts in Class II (Figure S3). As the DNA-binding motif of RAP2.3 is known (Franco-Zorrilla et al. 2014), we also determined what genes found in Clusters I and II, included the RAP2.3 DNA-binding elements in their promoters (3000 bases upstream of the transcription start site; 65 and 141 were found in Class I and Class II, respectively). For these genes we generated heatmaps of significant changes in transcript expression to determine under what specific stress conditions they are altered. As shown in Figures 4B and S4, Class I transcripts, encoded by genes with a RAP2.3 DNA-binding element, included several interesting TFs such as MYB51, WRKY6, and SIGMA3, proteins involved in redox/ROS regulation, such as Cupper/Zinc superoxide dismutase 2 and glutathione S-transferase, as well as proteins involved in DNA repair, MAPK cascades, osmotic stress, and different kinases and phosphatases. Class II transcripts, encoded by genes with a RAP2.3 DNA-binding element, included different ribosomal proteins, phosphatases, kinases, metal-binding proteins, calcium sensing proteins, development-related proteins, the TF WRKY33, and other transcripts (Figures 4 and S5; Tables S4-S6).

**Figure 4.**
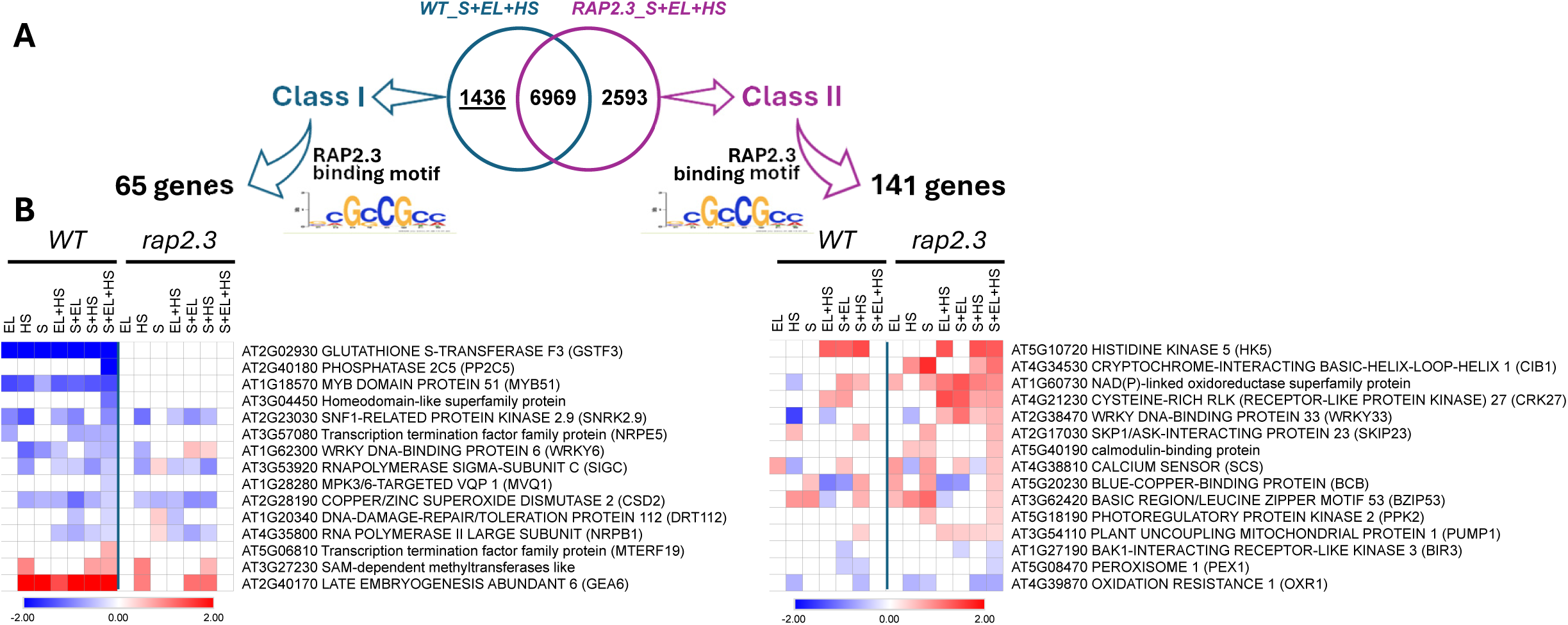
Identification of transcripts encoded by genes with a RAP2.3 binding element in their promoter during multifactorial stress combination (MFSC). **A.** A Venn diagram showing the definition of Class I and Class II transcripts in wild type (WT) and *rap2.3_1* seedlings grown under control conditions (CT) or subjected to a MFSC of salt (S), excess light (EL), and/or heat stress (HS). **B.** Heatmaps for the expression of selected Class I and Class II transcripts in WT and *rap2.3_1* seedlings subjected to S, EL, and/or HS in all possible combinations (heatmaps for all transcripts are shown in Figures S4 and S5). Abbreviations: EL, excess light; HS, heat stress; S, salt; WT, wild type Col0.

### 3.5. MYB51 and SIGMA3 are essential for plant survival under conditions of MFSC

To functionally characterize some of the transcripts identified by our transcriptomics analysis (Figures 2-4), we obtained two independent homozygous mutant alleles for 10 different genes found in Clusters I and II (Figure 5A, Table S2) and determined their survival under conditions of CT, S, EL, HS, and S+EL+HS (Figure S6). This analysis revealed that *myb51* (an R2R3-MYB TF that regulates glucosinolate metabolism in response to different stimuli) and *sig3* (a nuclear-encoded transcriptional regulator that directs the plastid-encoded RNA polymerase complex to specific promoters) mutants are sensitive to the 3-stress MFSC, but not to each of the individual stress conditions. In addition, as shown in Figure S6, two independent homozygous mutant for *mvq1* (MPK3/6-Targeted VQ motif-containing proteins 1; AT1G28280; a plant-specific transcriptional regulator that interacts with WRKY TFs to regulate plant growth, development, and responses to biotic/abiotic stressors; Kim et al., 2013; Pecher et al., 2014), and *phl13* (a homeodomain-like superfamily protein/PHR 13, AT3G04450; involved in phosphate starvation response; Huang et al., 2018), were also found to be sensitive to the 3-stress MFSC, but not to each of the individual stress conditions. In the current study we elected to focus on MYB51 and SIGMA 3 and will address the function of MVQ1 and PHL13 mutants in future studies.

**Figure 5.**
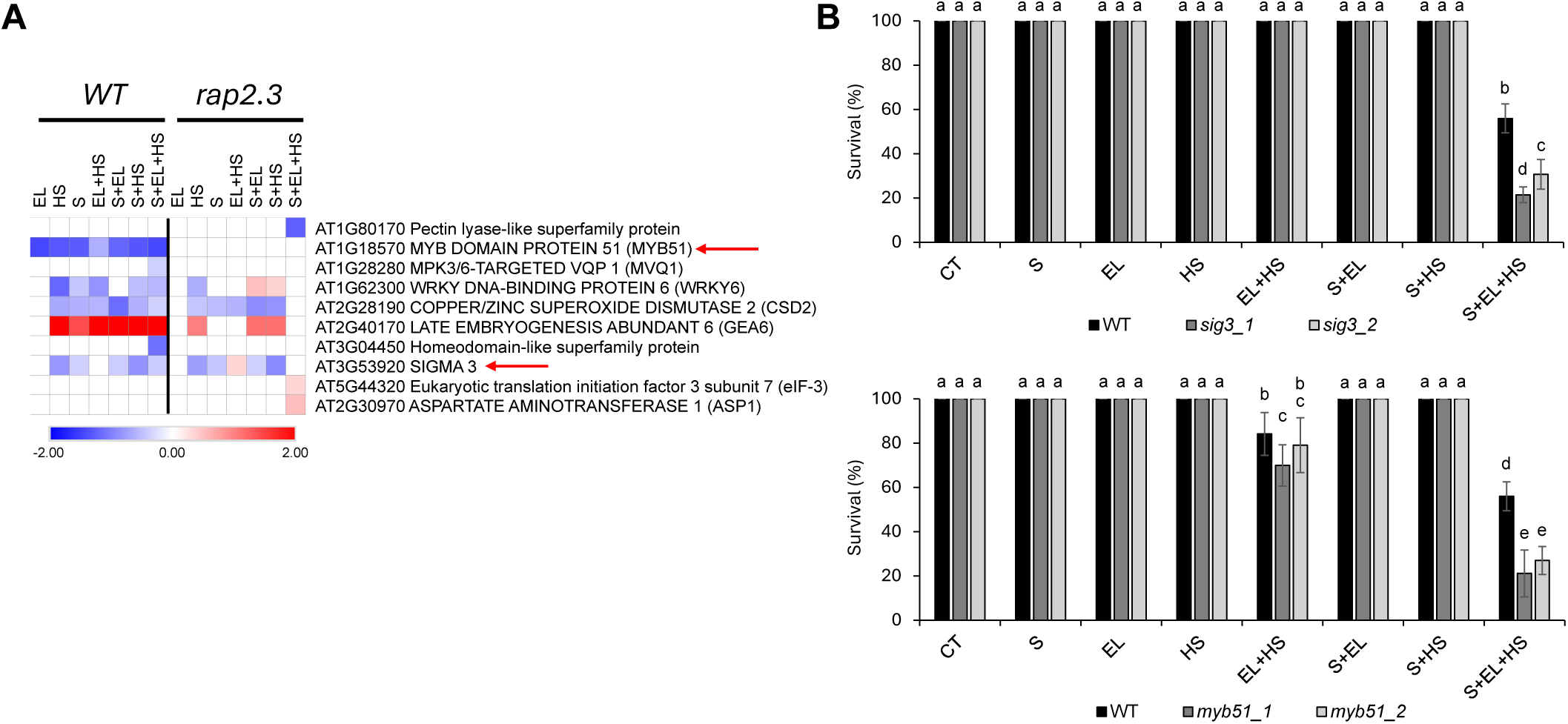
Survival of *myb51* and *sig3* mutants subjected to multifactorial stress combination. **A.** Heatmap of the expression pattern of all transcripts used for the genetic testing of survival in response to a multifactorial stress combination (MFSC) of salt (S), excess light (EL), and/or heat stress (HS) in all possible combinations (shown in Figures S4 and S5). **B.** Survival of *sig3* (*sig3_1*, SALK_009166 and *sig3_2*, CS879277; top) and *myb51* (*myb51_1*, CS306634 and *myb51_2*, CS924242; bottom) mutants grown under controlled growth conditions (CT), or subjected to a MFSC of S, EL, and/or HS, in all possible combinations. Two-way ANOVA followed by Fisher’s LSD post hoc test was used to determine significant differences between means (different letters denote statistically significant differences; *P* ≤ 0.05). Abbreviations: EL, excess light; HS, heat stress; S, salt; WT, wild type Col0.

To determine whether the sensitivity of *myb51* and *sig3* mutants is specific for the 3-stress MFSC, we tested the survival of *myb51* and *sig3* mutants to the application of S, EL and HS in all possible combinations. As shown in Figure 5B, like *bhlh35* (Sinha et al. 2025c) and *rap2.3* (Figure 1) mutants, *myb51* and *sig3* mutants were specifically susceptible to the S+EL+HS MFSC. Thus, a putative hierarchical order of expression regulation during the S+EL+HS MFSC could be: bHLH35 regulates RAP2.3, that in turn is required for MYB51 and SIGMA3 expression, and all four proteins are specifically required for survival under these MFSC conditions (Figures 1, 5B; Sinha et al. 2025c).

### 3.6. Gene regulatory network analysis of MYB51 reveals a potential role for JA signaling in plant responses to MFSC

MYB51 was previously found to be involved in regulating glucosinolate (GSL) metabolism in response to different pathogens and other signals, as well as linked to oxidative and osmotic stress responses in Arabidopsis (Gigolashvili et al. 2007; Dubois et al. 2013; Frerigmann and Gigolashvili, 2014; Zhao et al. 2017; De Clercq et al. 2021). We therefore decided to focus our analysis on this TF and conducted a GRN analysis of its function using the RNA-seq data we obtained for WT plants subjected to CT or S+EL+HS. As shown in Figure 6A, our GRN analysis of MYB51 identified 4 clusters (Table S9). Of these (due to transcript number in each cluster required for statistical analysis), we could perform GO analysis on cluster 6 (Figure 6B, Table S10) and cluster 3 (Figure S7, Table S10). Cluster 6 was found to be enriched in multiple transcripts associated with stress hormones and signal transduction processes, including JA signaling. In addition, this cluster included transcripts involved in GSL biosynthesis (Figure 6B, Table S10). Cluster 3, on the other hand, was enriched in temperature response, auxin signaling, metabolic pathways and other more ‘general function’-associated transcripts (Figure S7, Table S10). In accordance with these findings, DNA-binding elements for MYB51 and/or RAP2.3 were found in the promoters of multiple JA biosynthesis genes (Figure 7A). In addition, compared to WT, the expression of many transcripts involved in JA biosynthesis and function, including *JAR1* (involved in converting JA into its biologically active form, JA-Ile), *ACX3*, *OPR3*, *OPLC1*, and *CYP94B1* (involved in JA-Ile removal), was upregulated in the *rap2.3_1* mutant during the MFSC of S+EL+HS (associated with reduced survival of the *rap2.3_1* mutant to this stress combination; Figures 1A, 7A). JA was previously found to play an important role in the response of mature Arabidopsis and tomato plants to a combination of EL+HS (Balfagón et al. 2019; Pascual et al. 2023; He et al. 2025), suggesting that it could also be involved in plant survival to the MFSC of S+EL+HS. We therefore tested the survival of mutants deficient in JA perception (*coi1;* Stotz et al. 2011; Howe et al. 2018) or biosynthesis (*jar1;* Staswick and Tiryaki, 2004; Howe et al. 2018), subjected to S, EL, and HS in all possible combinations. As shown in Figure 7B, compared to WT, *jar1* and *coi1* mutants were significantly more resistant to the S+EL+HS combination. This finding suggests that the JA pathway could play an antagonistic role in plant survival to the combination of S+EL+HS, and that its suppression in WT during S+EL+HS, compared to *rap2.3_1* (Figure 7B), promotes survival under these conditions. Further studies are needed, however, to determine whether RAP2.3, through MYB51, suppresses JA signaling to promote survival under conditions of S+EL+HS.

**Figure 6.**
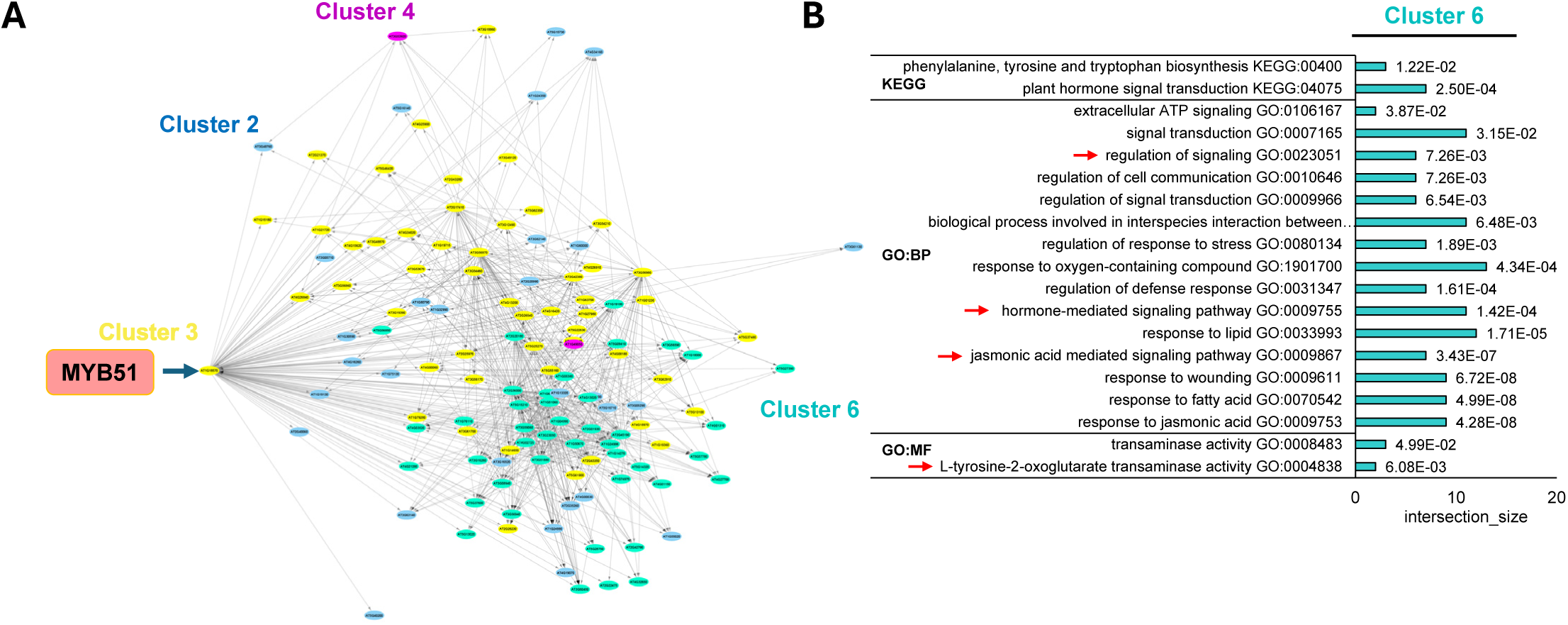
Gene regulatory network analysis of *MYB51* during multifactorial stress combination. **A.** Gene regulatory network (GRN; Differential inference analysis of expression; DIANE) analysis of *MYB51* during a multifactorial stress combination (MFSC) of salt (S), excess light (EL), and heat stress (HS). **B.** Gene ontology (GO) analysis of cluster 4 identified as associated with *MYB51* function during a MFSC of S+EL+HS. Abbreviations: BP, biological process; GO, gene ontology; MF, molecular function; KEGG, Kyoto Encyclopedia of Genes and Genomes.

**Figure 7.**
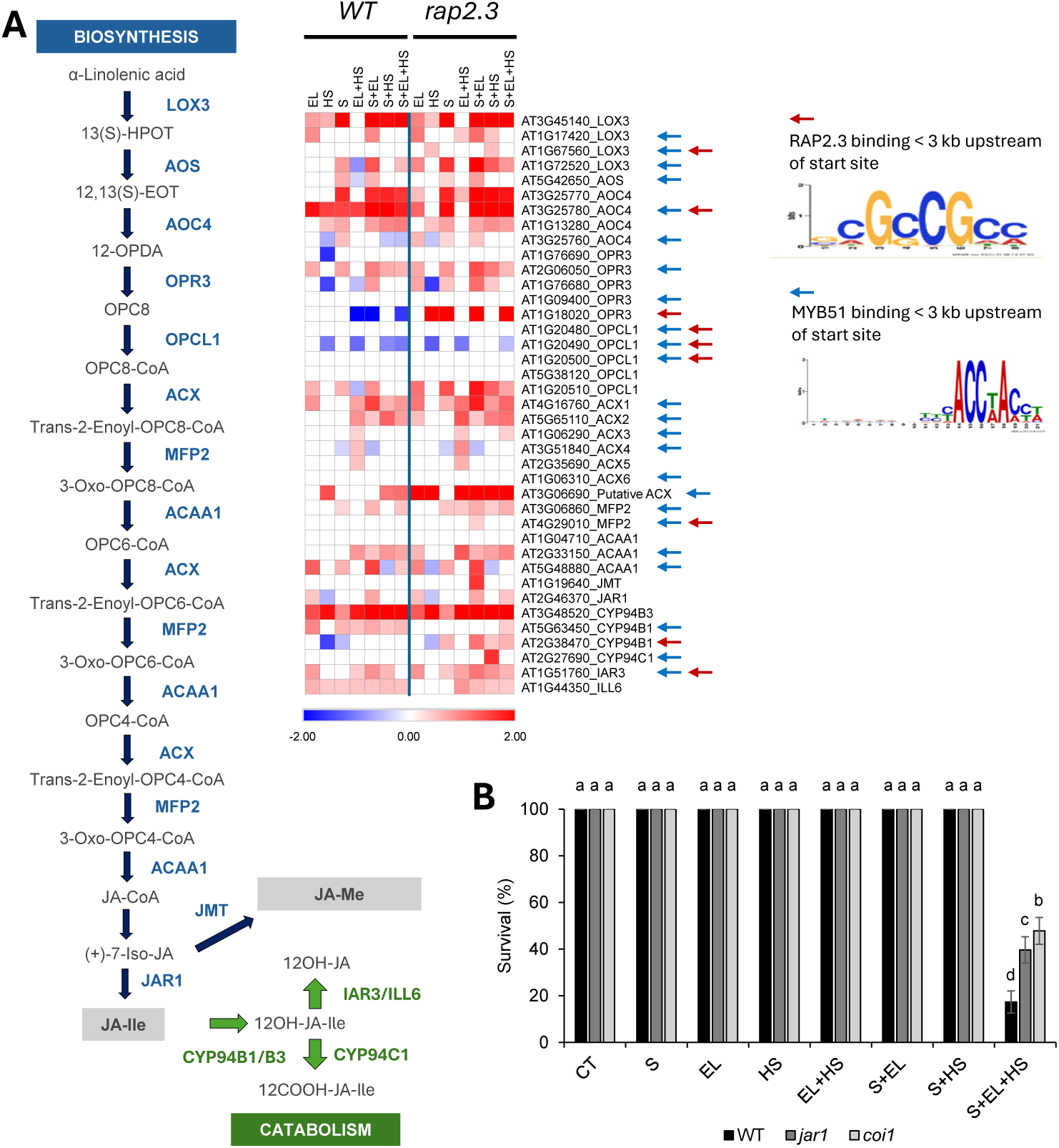
Involvement of jasmonic acid (JA) in plant responses to multifactorial stress combination. **A.** A schematic model of JA biosynthesis and catabolism pathways (left) and a heatmap for the expression of transcripts involved in JA metabolism and catabolism in wild type (WT) and *rap2.3_1* seedlings grown under control conditions (CT) or subjected to a multifactorial stress combination (MFSC) of salt (S), excess light (EL), and heat stress (HS). Arrows next to the heatmap indicate transcripts encoded by genes that contain MYB51 and/or RAP2.3 DNA binding elements in their promoters. **B.** Survival of *jar 1* (SALK_030821C) and *coi1* (SALK_095916C) mutants grown under controlled CT, S, EL, and/or HS, in all possible combinations. Two-way ANOVA followed by Fisher’s LSD post hoc test was used to determine significant differences between means (different letters denote statistically significant differences; *P* ≤ 0.05). Abbreviations: EL, excess light; HS, heat stress; S, salt; WT, wild type Col0; LOX3, lipoxygenase 3; 13 (S)-HPOT, 13-Hydroperoxyoctadeca-9,11,15-trienoic acid; 12,13(S)-EOT, (9Z,15Z)-(13S)-12,13-Epoxyoctadeca-9,11,15-trienoic acid; AOS, allene oxide synthase; 12-OPDA, (9S,13S)-12-Oxophytodienoate; AOC4, ALLENE OXIDE CYCLASE 4; OPR3, 12-oxophytodienoic acid reductase; OPC8, (9S,13S)-10,11-Dihydro-12-oxo-15-phytoenoate; OPC1, OPC-8:0 CoA ligase 1; ACX1, Acyl-CoA oxidase 1; MFP2, enoyl-CoA hydratase/3-hydroxyacyl-CoA dehydrogenase; ACAA1, acetyl-CoA acyltransferase 1; JMT, JA carboxyl methyltransferase; JAR1, JA-amide synthetase 1; Cytochromes P450 CYP94C1 and CYP94B1/B3; IAR3, IAA-alanine resistant 3; ILL6, IAA-amino acid hydrolase ILR1-like 6.

## 4. DISCUSSION

RAP2.3 is an ethylene-response DNA binding protein belonging to the APETALA 2/Ethylene Response Factor (AP2/ERF) group VII of the ERF TF family (Nakano et al. 2006). This key transcriptional regulator is involved in modulating different plant responses to anaerobic stress and pathogen infection, as well as in controlling stomatal aperture and regulating different developmental processes (Chen et al. 2014; Papdi et al. 2015; Gibbs et al. 2015; Liu et al. 2018; León et al. 2020; Li et al. 2022). RAP2.3 also plays a key role in NO, abscisic acid (ABA), JA, and Gibberellin (GA) signaling (Gibbs et al. 2014; la Rosa et al. 2014; León et al. 2020). In addition to being transcriptionally regulated, the levels of RAP2.3 are post-translationally regulated by the N-degron–E3 ubiquitin ligases pathway that targets it for degradation unless it is turned off, for example under anaerobic conditions (Gibbs et al. 2015). Our findings that the basal levels of the *RAP2.3* transcripts are elevated in the *rap2.3* mutants and that in the *rap2.3* mutants *RAP2.3* expression is higher than in WT in response to the MFSC of S+EL+HS, correlating with *rap2.3* sensitivity to S+EL+HS (Figures 1, 2), could suggest that the high expression levels of *RAP2.3* in the *rap2.3* mutants results in RAP2.3 accumulation that in turn cause enhanced sensitivity of plants to S+EL+HS. This possibility is supported by our findings that overexpression of *RAP2.3* in WT plants (driven by the 35S-CaMV promoter) also causes enhanced sensitivity of plants to S+EL+HS conditions (Figure 1D), and that it alters NO levels (Figure 1F). However, it is presently unknown whether the N-degron–E3 ubiquitin ligases pathway is turned on or off during conditions of S+EL+HS in cells. If indeed this pathway is turned off (Figure S1), or overwhelmed by the overaccumulation of the *RAP2.3* transcripts (Figure 1C-F), then the accumulation of RAP2.3 protein in cells could function as a negative regulator of pathways required for S+EL+HS survival, or as a positive regulator of pathways that conflict with survival under the S+EL+HS MFSC. Further studies are needed to resolve this question.

Our study identified two transcriptional regulators (SIGMA3 and MYB51) that were expressed in WT, but not in the *rap2.3* mutant, in response to the S+EL+HS MFSC (Figure 5A), and mutants of these transcriptional regulators displayed an S+EL+HS-specific reduced survival phenotype, similar to that of the *rap2.3* (Figures 5B and 1A) and *bhlh35* (Sinha et al., 2025c) mutants. Although both transcriptional regulators contain RAP2.3 DNA-binding elements in their promoter (Figure S4) and their expression is dependent on *RAP2.3* overexpression in the *rap2.3* mutants (Figures 4, 5), further studies are needed to reveal whether RAP2.3 binds to the promoters of SIGMA3 and MYB51 (*e.g.,* electrophoresis mobility shift assays, DAP-/ChIP-Seq, and luciferase reporter assays).

SIGMA3 is one of six nuclear-encoded Arabidopsis sigma factors that direct the plastid-encoded RNA polymerase complex to specific plastid promoters (Zghidi et al. 2007; Chi et al. 2015). SIGMA3 is primarily involved in transcribing the *psbN* mRNA, encoding a crucial subunit of photosystem II required for efficient assembly and repair of PSII following photoinhibition (Zghidi et al. 2007; Chi et al. 2015). Our findings that *sig3* mutants are impaired in survival specifically under the MFSC of S+EL+HS (Figure 5B), could suggest that PSII repair and function are required for survival under MFSC conditions. This finding highlights the role of the chloroplast in maintaining survival and function under conditions of MFSC and adds to the previously identified roles for cytosolic and nuclear processes in promoting resilience to MFSC in plants (Zandalinas et al. 2021b; Pascual et al. 2025a; Sinha et al. 2025c). Beyond its specific role in *psbN* transcription, SIGMA3 may point to plastid gene expression as a critical bottleneck for acclimation to S+EL+HS. Excess light and heat can increase excitation pressure on PSII and enhance the need for rapid PSII repair, whereas salinity can further constrain photosynthetic performance through osmotic and ionic stress. Under the MFSC of S+EL+HS, impaired SIGMA3-dependent plastid transcription could reduce the capacity of chloroplasts to maintain photosynthetic homeostasis. This interpretation is consistent with the specific role of SIGMA3 in plastid *psbN* transcription (Zghidi et al., 2007), the requirement of PsbN for PSII reaction center assembly and recovery from photoinhibition (Torabi et al., 2014), and the broader function of plastid sigma factors in coordinating plastid transcriptional responses with nuclear-encoded regulatory inputs (Chi et al., 2015). In future studies, it will be important to determine how disruption of SIGMA3 function impairs plant resilience to MFSC.

MYB51 is a R2R3-MYB TF that regulates indolic glucosinolates (IGs) biosynthesis in response to different biotic stimuli (Frerigmann and Gigolashvili, 2014; Frerigmann et al. 2016). GSLs are secondary metabolites that play a key role in plant defense against pathogens and herbivores (Halkier and Gershenzon, 2006). Expression of MYB51 was also reported to increase in Arabidopsis in response to different abiotic stresses, including osmotic and oxidative stress (Burow, 2016; Zhao et al. 2017; De Clercq et al. 2021), and MYB51 expression was associated with Ethylene Response Factor 6 (ERF6; belonging to the same AP2/ERF family as RAP2.3; Dubois et al. 2013) and WRKY33 (identified in Class II RAP2.3 transcripts; Figure S5; Barco and Clay, 2020) function. The identification of MYB51 as a TF that functions downstream of RAP2.3 and, like bHLH35, RAP2.3, and SIGMA3, is specifically required for plant survival under conditions of S+EL+HS (Figure 5B), could suggests that survival under conditions of S+EL+HS MFSC requires tolerance to oxidative stress (as suggested by Sinha et al. 2025c) and/or IGs biosynthesis. In addition, MYB51 could be involved in suppressing the JA pathway to promote survival under conditions of S+EL+HS (Figures 6, 7). An additional possibility is that MYB51 contributes to the prioritization of stress responses under S+EL+HS by balancing defense metabolism with acclimation-related processes. Although indolic glucosinolate- and JA-associated pathways are central components of Arabidopsis defense and stress signaling (Halkier and Gershenzon, 2006; Frerigmann et al., 2016; Howe et al., 2018), their excessive activation during S+EL+HS could potentially impose metabolic costs or divert resources from processes required for survival, including photosynthetic repair, redox homeostasis and growth maintenance (Mittler, 2006; Zandalinas and Mittler, 2022; Rivero et al., 2022). Thus, a RAP2.3–MYB51-dependent regulation of gene expression may help coordinate the magnitude and timing of defense-associated responses so that they remain compatible with MFSC acclimation. This interpretation is consistent with the enhanced survival of *coi1* and *jar1* mutants under S+EL+HS, while preserving the idea that JA function is highly context dependent during combined stress responses (Balfagón et al., 2019; Howe et al., 2018; Pascual et al., 2023).

Our findings that activation of the JA pathway could be associated with the enhanced sensitivity of the *rap2.3_1* mutant to a MFSC of S+EL+HS (Figure 7), coupled with RAP2.3 proposed role in ethylene signaling (Gasch et al. 2016; Liu et al. 2018; Kim et al. 2018; Sinha et al. 2025c), could suggest an antagonistic interaction between ethylene and JA signaling during MFSC. Ethylene and JA signaling were found to have synergistic or antagonistic interactions, depending on stress type and developmental stage (Tuominen et al. 2004; Song et al. 2014; Zhu, 2014; He et al. 2017; Wang et al. 2025). As we previously reported, ethylene signaling during MFSC could be mediated by RAP2.3 and mutants impaired in ethylene signaling display altered survival under conditions of S+EL+HS, as well as EL+HS (Sinha et al. 2025c). As the function of RAP2.3 is altered in the *rap2.3_1* mutant, the ethylene signaling pathway in this mutant may not suppress JA signaling resulting in the elevated expression of *JAR1*, *ACX3*, *OPR3*, *OPLC1*, and *CYP94B1* in the *rap2.3_1* mutant during MFSC, compared to WT (Figure 7). Thus, the altered expression of RAP2.3 in the *rap2.3* mutants may suppress the ethylene pathway resulting in the activation of the JA pathway during MFSC (in *rap2.3* mutants), that results in their reduced survival. Further studies are needed to address this intriguing possibility, including measurements of ethylene and JA levels and side-by-side analysis of ethylene and JA signaling mutants subjected to MFSC.

## 5. SUMMARY

Our findings reveal that RAP2.3 plays an essential role in plant tolerance to a MFSC of S+EL+HS, and that this TF is required for the expression of SIGMA3, required for PSII function specifically during S+EL+HS, as well as for the expression of MYB51, potentially involved in JA and IGs/GSLs signaling during MFSC. These findings could define a facet of plant metabolism and function under conditions of MFSC, which is different from that defined for bHLH35 (Sinha et al. 2025c). Thus, in addition to flavonoids and ROS metabolism controlled by bHLH35 (Sinha et al. 2025c), photosynthesis, controlled by SIGMA3, and/or JA and/or IGs/GSLs signaling, potentially controlled by MYB51 (both regulated by RAP2.3), are also required for plant survival under conditions of MFSC. In future studies it would be interesting to examine whether RAP2.3 functions as a suppressor of pathways required for tolerance to MFSC, or as an activator of pathways that conflict with plant survival to MFSC (*e.g.,* JA; Figure 7). Thus, RAP2.3 could suppress the expression of an important positive regulator of plant responses to MFSC (*e.g.,* SIGMA3), thereby resulting in enhanced sensitivity of plants to S+EL+HS. Alternatively, it could suppress or enhance the expression of pathways that are antagonistic to MFSC survival (*e.g.,* JA).

## ACKNOWLEDGMENTS

We thank the Arabidopsis Biological Resource Center (https://abrc.osu.edu/) for seeds of Arabidopsis mutants used in this study.

## AUTHOR CONTRIBUTIONS

Conceptualization: RM, SIZ, RS; Methodology: RS, MAPV, DM, ZL, LSP, AB; Investigation: RM, SIZ, RS, MAPV, TJ, RKA; Visualization: RS, MAPV, AB, ZL; Funding acquisition: RM, TJ; Project administration: RM; Supervision: RM, TJ; Writing – original draft: RM, RS; Writing – review & editing: All authors.

## DATA AND MATERIALS AVAILABILITY

All materials used in the analysis are available upon request from the Corresponding Author. All data are available in the main text or the supplementary materials. RNA-Seq data is available at Gene Expression Omnibus (GEO) under accession GSE330879.

## SUPPLEMENTARY MATERIALS

### SUPPLEMENTARY FIGURE LEGENDS

**Figure S1.**
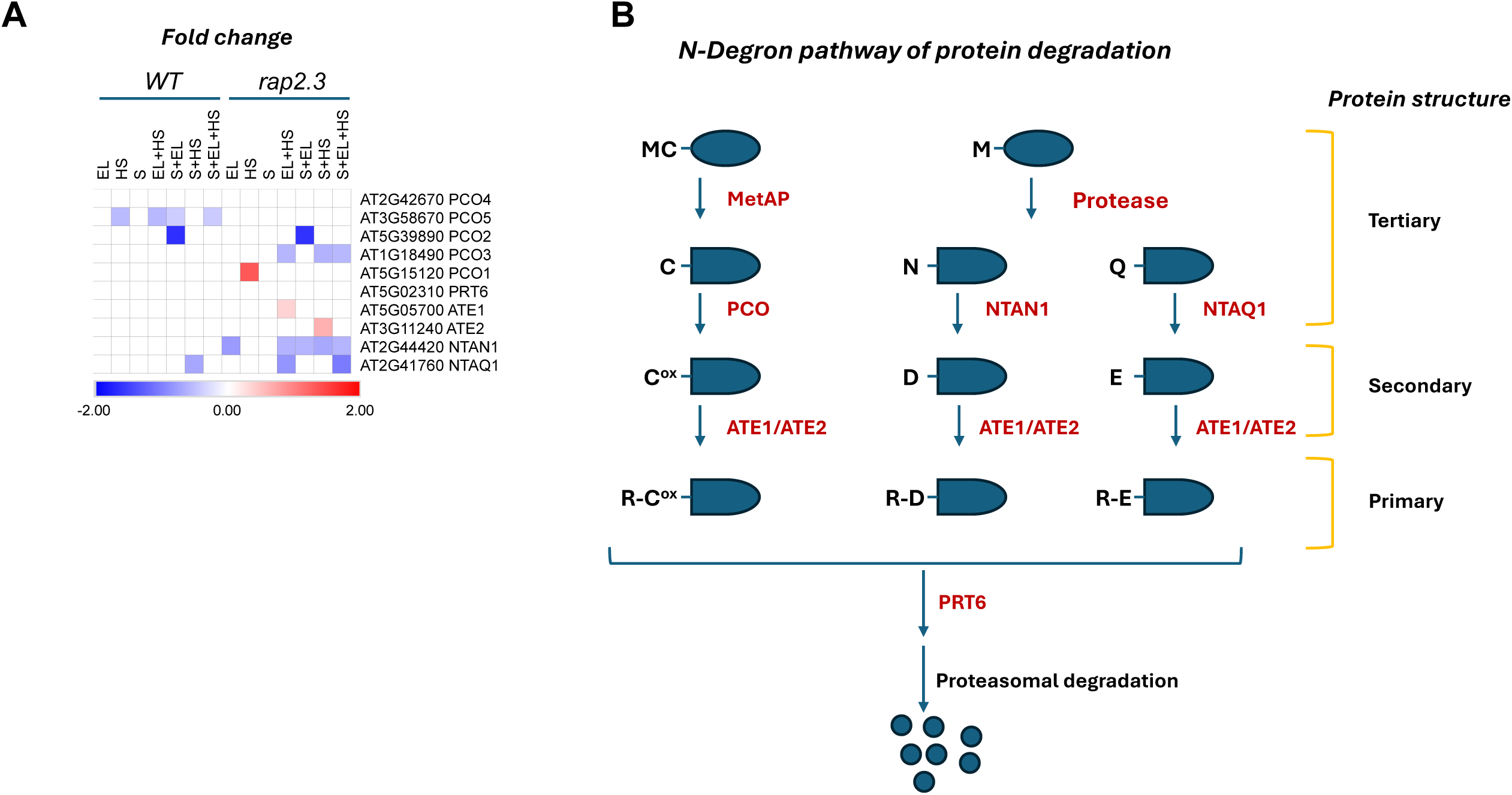
Expression of transcripts encoding enzymes involved in the N-degron pathway of protein degradation under MFSC. **A.** Heatmap for the relative change in the expression of transcripts involved in the N-degron pathway of protein degradation under conditions of MFSC. **B.** Illustration of the N-degron pathway of protein degradation. Abbreviations: ATE, Arginyl-Transfer RNA (tRNA):Protein Arginyl Transferase; EL, excess light; HS, heat stress; S, salt; WT, wild type Col0; PCO, Plant Cysteine Oxidase; PRT, Proteolysis; NTAN1, Asn-specific N-terminal amidase; NTAQ1, Gln-specific N-terminal amidase; MetAP, Met aminopeptidases; M, Methionine; C, Cysteine; C^ox^, Oxidized cysteine; D, Aspartic acid; E, Glutamic acid; R, Arginine; N, Asparagine; Q, Glutamine. In support of Figure 1.

**Figure S2.**
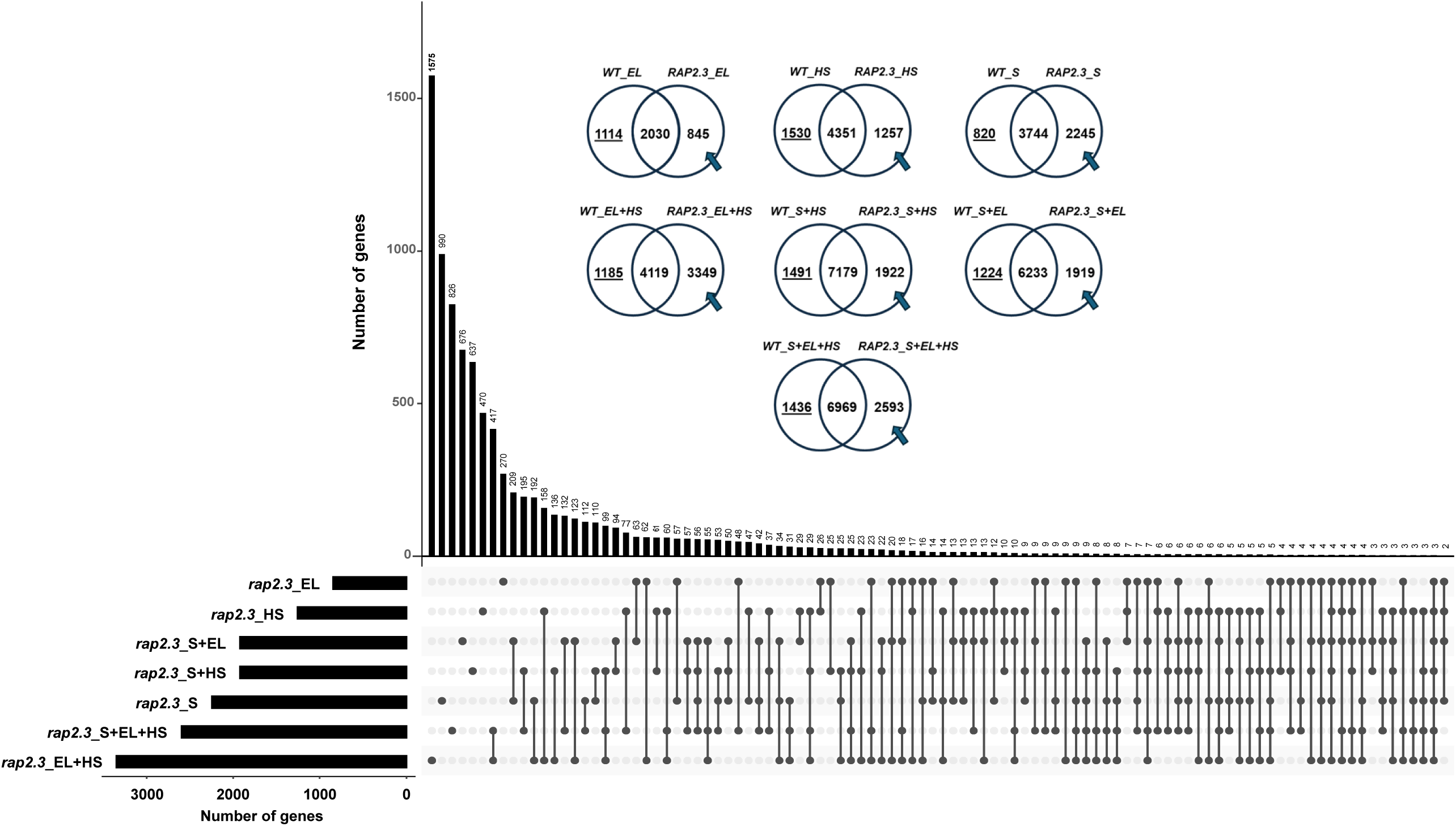
Transcriptomics analysis of WT and *rap2.3_1* seedlings subjected to salt (S), excess light (EL) and heat stresses (HS) in all possible combinations. Venn diagrams showing the overlap between transcripts significantly altered in their expression in WT or the *rap2.3_1* mutant in response to all stress treatments compared to CT. Arrows indicate transcripts with significantly altered expression in *rap2.3_1*, but not in WT. The UpSet plot shows the overlap between all transcripts indicated with an arrow in the Venn diagrams. Negative binomial Wald test followed by Benjamini-Hochberg correction (P < 0.05) was used for statistical significance between transcript expressions. Abbreviations: CT, control; EL, excess light; HS, heat stress; S, salt; WT, wild type Col0. In support of Figure 2.

**Figure S3.**
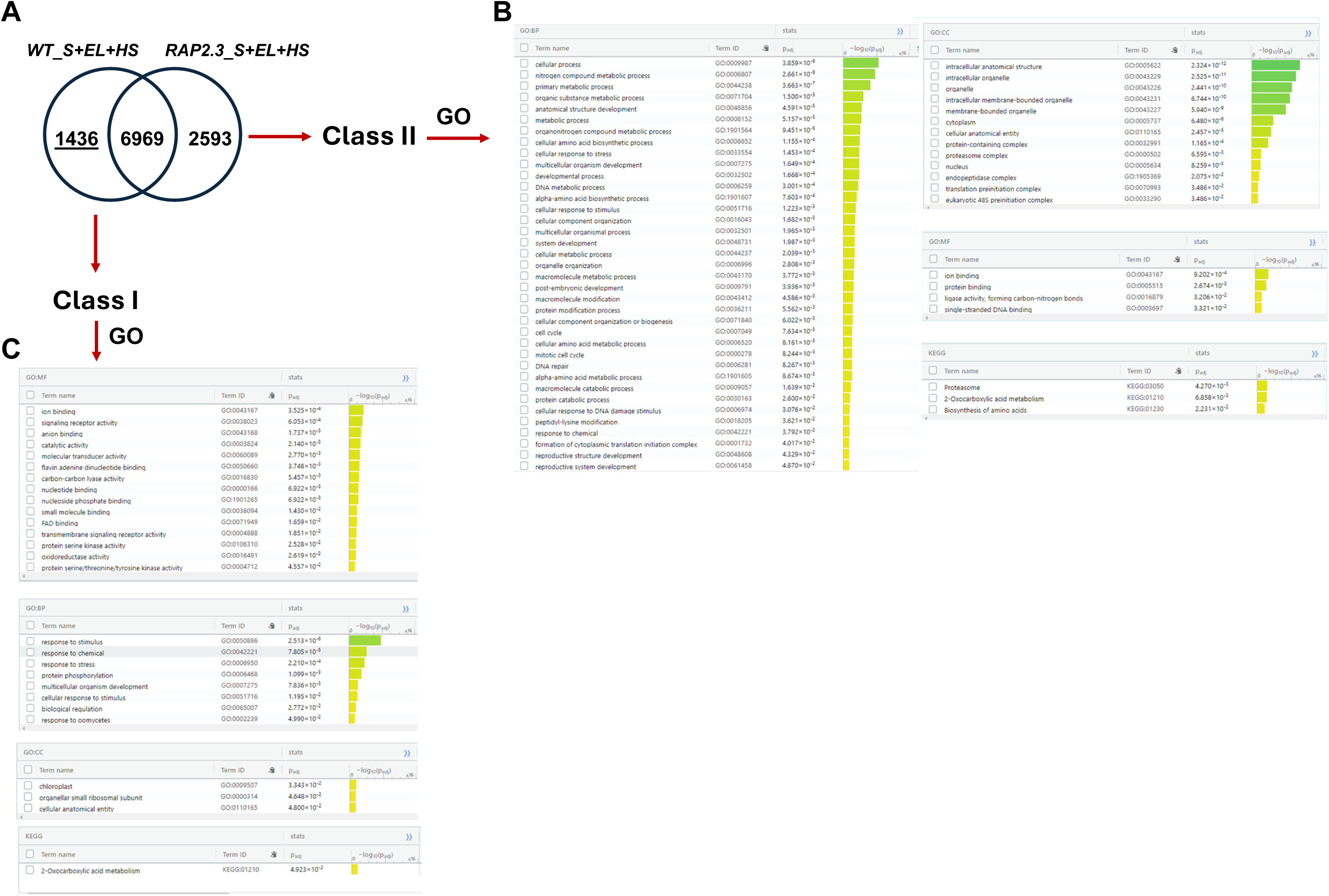
Gene ontology analysis of transcripts unique to WT or *rap2.3* during multifactorial stress combination (MFSC). **A.** A Venn diagram showing the definition of Class I and Class II transcripts in wild type (WT) and *rap2.3_1* seedlings grown under control conditions (CT) or subjected to a MFSC of salt (S), excess light (EL), and/or heat stress (HS). **B.** Gene ontology analysis of transcripts unique to *rap2.3* during MFSC of S+EL+HS. **C.** Gene ontology analysis of transcripts unique to WT during a MFSC of S+EL+HS. Abbreviations: BP, biological process; GO, gene ontology; MF, molecular function, KEGG, Kyoto Encyclopedia of Genes and Genomes. In support of Figure 4.

**Figure S4.**
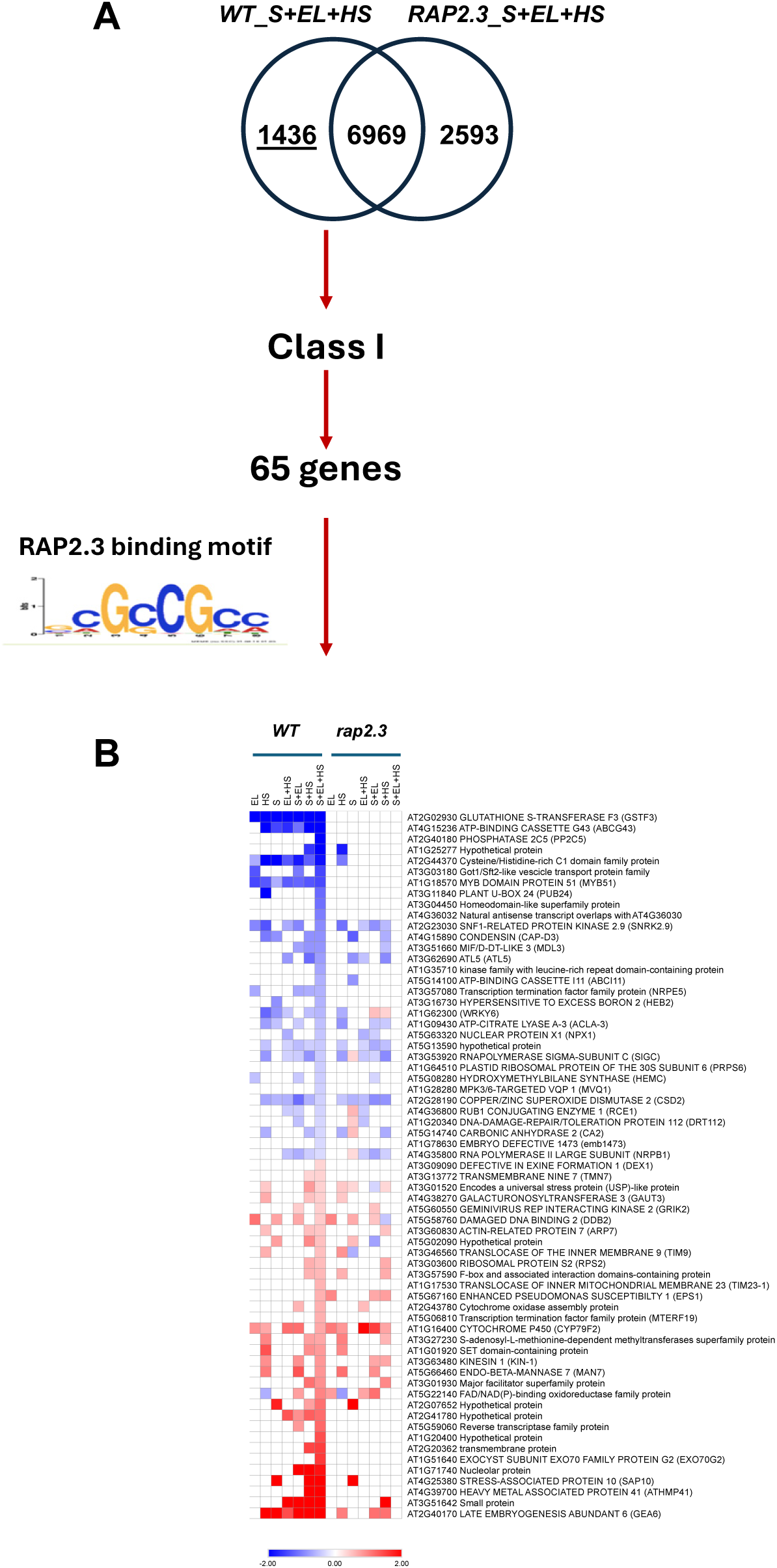
Identification of transcripts encoded by genes with a RAP2.3 binding element in their promoter during multifactorial stress combination (MFSC). **A.** A Venn diagram showing the definition of Class I and Class II transcripts in WT and *rap2.3_1* seedlings grown under control conditions (CT) or subjected to a MFSC of salt (S), excess light (EL), and/or heat stress (HS). **B.** Heatmap for the expression of Class I transcripts containing RAP2.3 binding motif in their promoter in WT seedlings subjected to S, EL, and/or HS in all possible combinations. Abbreviations: EL, excess light; HS, heat stress; S, salt; WT, wild type Col0. In support of Figure 4.

**Figure S5.**
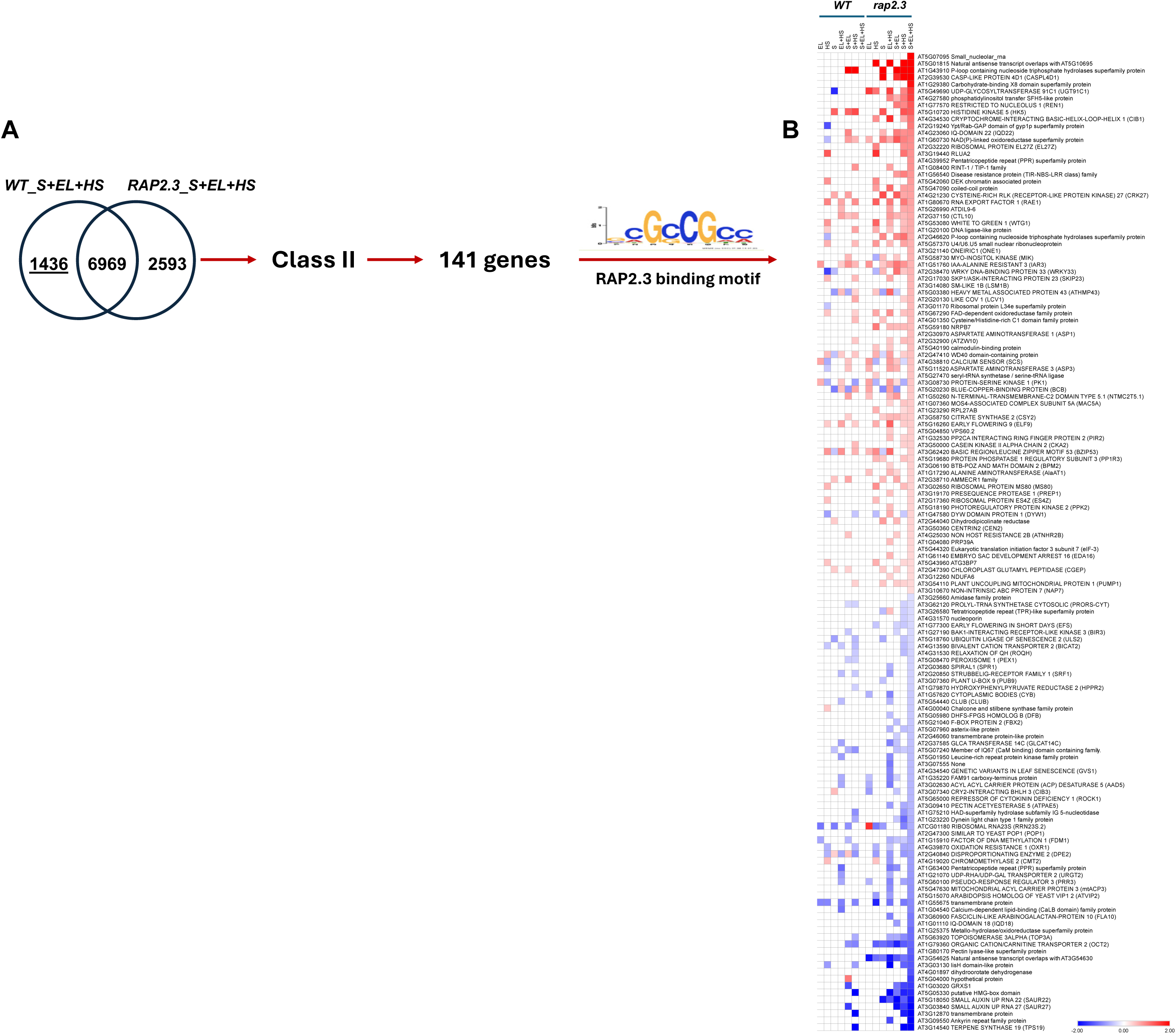
Identification of transcripts encoded by genes with a RAP2.3 binding element in their promoter during multifactorial stress combination (MFSC). **A.** A Venn diagram showing the definition of Class I and Class II transcripts in WT and *rap2.3_1* seedlings grown under control conditions (CT) or subjected to a MFSC of salt (S), excess light (EL), and/or heat stress (HS). **B.** Heatmap for the expression of Class II transcripts containing RAP2.3 binding motif in their promoter in *rap2.3_1* seedlings subjected to S, EL, and/or HS in all possible combinations. Abbreviations: EL, excess light; HS, heat stress; S, salt; WT, wild type Col0. In support of Figure 4.

**Figure S6.**
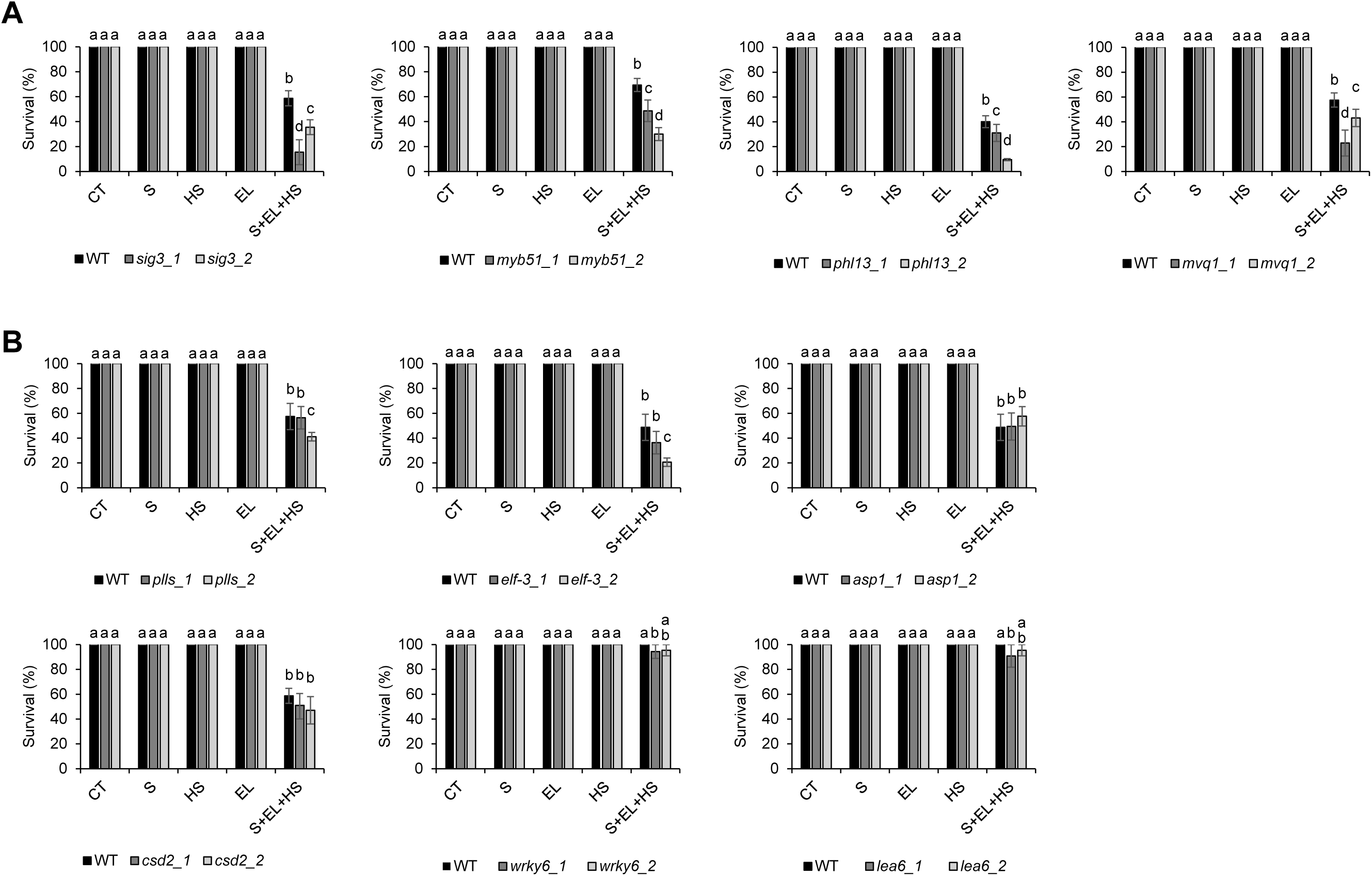
Survival of mutants subjected to multifactorial stress combination (MFSC). **A.** Survival of SIGMA 3 (*sig3_1*, SALK_009166C and *sig3_2*, SAIL_1289_G07/CS879277), MYB DOMAIN PROTEIN 51 (*myb51_1*, SALKseq_040923.3/CS924242 and *myb51_2*, GK-228B12/CS306634), Homeodomain-like superfamily protein (PHL13; *phl13_1*, SALK_203017C and *phl13_2*, SALK_036703), MPK3/6-Targeted VQ motif-containing proteins 1 (MVQ1; *mvq1_1*, SALK_091850 and *mvq1_2*, SALK_107266C) mutants grown under controlled growth conditions (CT), or subjected to a MFSC of salt (S), excess light (EL), and/or heat stress (HS), in all possible combinations. **B.** Survival of Pectin lyase-like superfamily protein (*plls_1*, SALK_040741 and *plls_2*, SALK_095457C), Eukaryotic translation initiation factor 3 subunit 7 (*elf-3_1*, SALK_085397C and *elf-3_2*, SALK_105336C), Aspartate Aminotransferase 1 (*asp1_1*, SALK_082821C and *asp1_2*, CS887402), Copper/Zinc Superoxide Dismutase 2 (*csd2_1*, CS882522 and *csd2_2*, SALK_041901.55.75), WRKY6 (*wrky6_1*, CS868868 and *wrky6_2*, SALK_012997) and Late Embryogenesis Abundant 6 (GEA6; *lea6_1*, SALK_041260C and *lea6_2*, SALK_112719C) mutants grown under controlled growth conditions (CT), or subjected to a MFSC of S, EL, and/or HS, in all possible combinations. Two-way ANOVA followed by Fisher’s LSD post hoc test was used to determine significant differences between means (different letters denote statistically significant differences; P ≤ 0.05). Abbreviations: EL, excess light; HS, heat stress; S, salt; WT, wild type Col0. In support of Figure 5.

**Figure S7.**
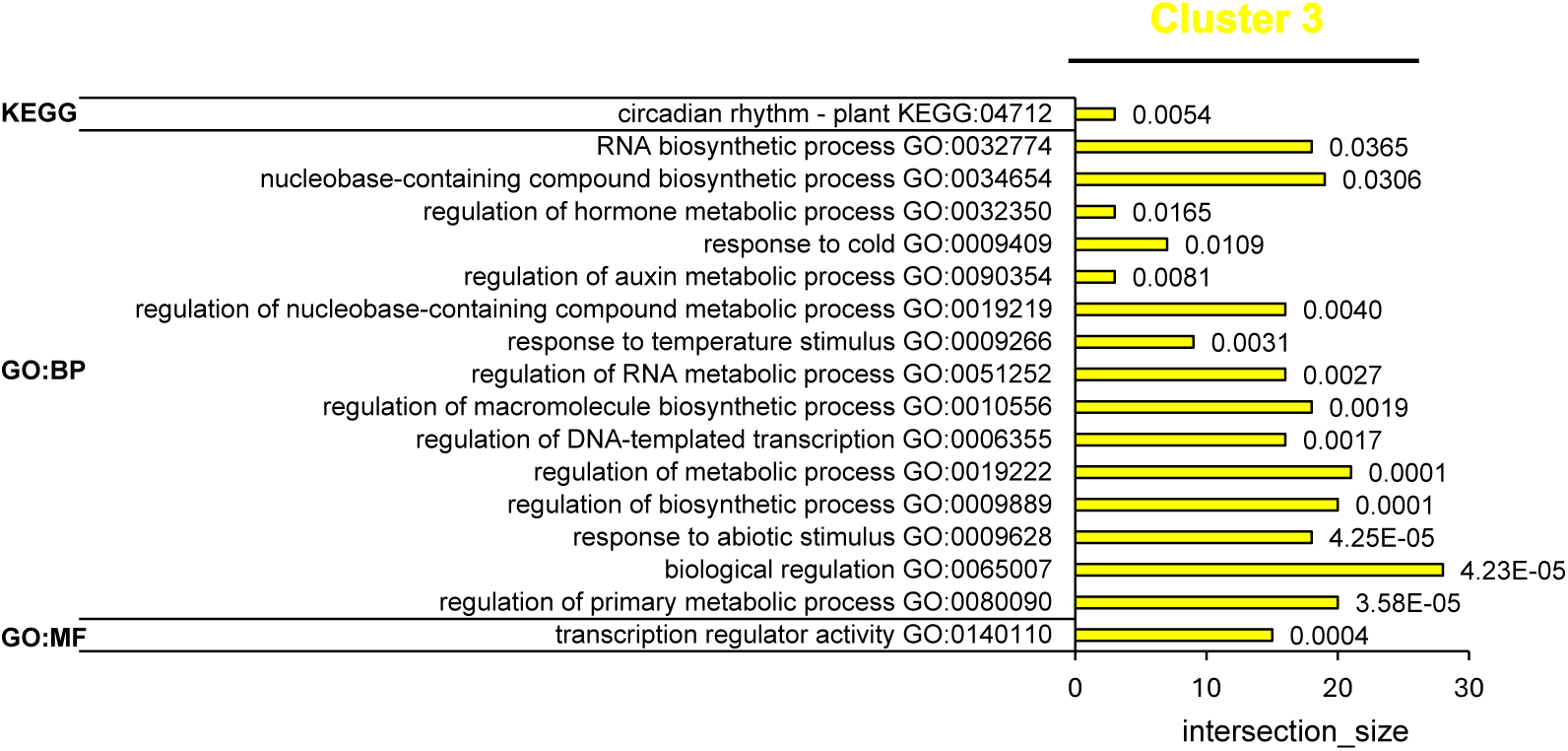
Gene regulatory network analysis of cluster 3 from MYB51 gene regulatory network analysis during multifactorial stress combination. Abbreviations: BP, biological process; GO, gene ontology; MF, molecular function, KEGG, Kyoto Encyclopedia of Genes and Genomes. In support of Figure 6.

### SUPPLEMENTARY TABLES

**Table S1:** Details of stress treatment used in the study.

**Table S2:** Details of *Arabidopsis thaliana* T-DNA insertion mutants used in the study.

**Table S3:** List of primers used in the study.

**Table S4:** Transcripts significantly altered in their expression in wild type Arabidopsis thaliana plants subjected to S, EL, and/or HS stresses (in all possible combinations compared to control conditions).

**Table S5:** Transcripts significantly altered in their expression in rap2.3 mutant Arabidopsis thaliana plants subjected to S, EL, and/or HS stresses (in all possible combinations compared to control conditions).

**Table S6:** Transcripts with overlapping or unique expression included in Figure 2B Venn diagrams.

**Table S7:** Description of genes in each cluster of the RAP2.3 gene regulatory network analysis (Figure 3A).

**Table S8:** Gene Ontology of genes in cluster 7 of the RAP2.3 gene regulatory network analysis (Figure 3B).

**Table S9:** Description of genes in each cluster of the MYB51 gene regulatory network analysis (Figure 6A).

**Table S10:** Gene Ontology of genes in cluster 3 and 6 of the MYB51 gene regulatory network analysis (Figures 6B and S7).

